# A Tumour-Specific Molecular Network Promotes Tumour Growth in *Drosophila* by Enforcing a JNK-YKI Feedforward Loop

**DOI:** 10.1101/2023.10.18.561369

**Authors:** Indrayani Waghmare, Karishma Gangwani, Arushi Rai, Amit Singh, Madhuri Kango-Singh

## Abstract

Cancer cells expand rapidly in response to altered intercellular and signalling interactions to achieve hallmarks of cancer. Impaired cell polarity combined with activated oncogenes is known to promote several hallmarks of cancer e.g., activating invasion by increased activity of Jun N-terminal kinase (JNK), and sustained proliferative signalling by increased activity of Hippo effector Yorkie (Yki). Thus, JNK, Yki, and their downstream transcription factors have emerged as synergistic drivers of tumour growth through pro-tumour signalling and intercellular interactions like cell-competition. However, little is known about the signals that converge onto JNK and Yki in tumour cells that enable the tumour cells to achieve hallmarks of cancer. Here, using mosaic models of cooperative oncogenesis (*Ras^V12^, scrib^-^*) in *Drosophila*, we show that *Ras^V12^, scrib^-^* tumour cells grow by activation of a previously unidentified network comprising Wingless (Wg), Dronc, JNK and Yki. We show that *Ras^V12^, scrib^-^* cells show increased Wg, Dronc, JNK, and Yki signalling, and all of these signals are required for the growth of *Ras^V12^, scrib^-^* tumours. We report that Wg and Dronc converge onto a JNK-Yki self-reinforcing positive feedback signal-amplification loop that promotes tumour growth. We found that Wg-Dronc-Yki-JNK molecular network is specifically activated in polarity-impaired tumour cells and not in normal cells where apical basal polarity is intact. Our findings suggest that identification of molecular networks may provide significant insights about the key biologically meaningful changes in signalling pathways, and paradoxical signals that promote Tumourigenesis.

## 1. Introduction

Over the past decade, genomic, transcriptomic and proteomic data from several cancers has revealed the genetic alterations and changes in gene expression and epigenetic modifications linked to several cancers (1–4). These approaches have revealed the extensive differences between normal and cancer cells, and the effects of cancer on multiple genes and signalling pathways(3). Although these studies have provided valuable information about cancer cells, their origin, oncogenic processes and signalling pathways, efforts now need to be directed at further characterizing tumours. For example, which proteins are expressed in the same cancer cell, and are likely to physically interact? Which signalling pathways play decisive roles in promoting cell proliferation, survival and metastasis in the different stages of the cancer? Do intricate distribution or clustering of ligand-receptors affect cell-cell interactions(5) For this, complementary approaches with *in-vivo* models that allow manipulation of multiple genes (which represent the key driver mutations of the cancer) are required.

*Drosophila*, the common fruit fly, represents an efficient model organism to modulate the expression of multiple genes to study the signalling crosstalk, which promotes tumour growth and progression (6,7). Models of cooperative oncogenesis in *Drosophila,* such as clonal tumours induced by activation of oncogenic Ras (*Ras^V12^*) in polarity deficient *scribble (scrib)* [*Ras^V12^ scrib^-^*], or *lethal giant larva (lgl)* or *discs large (dlg)* mutant cells, show classic hallmarks of aggressive cancer growth exemplified by increased proliferation rate, reduced apoptosis and differentiation, and metastasis (8–11). These models have been instrumental in establishing the links between deregulation of cell polarity, and increased signalling from Jun N-terminal Kinase (JNK) and Yorkie (Yki), effectors of two key signalling pathways that regulate cell proliferation and apoptosis (12–16). However, the upstream mechanisms by which JNK and Yki activation promote aggressive metastatic growth during oncogenic cooperation remain poorly defined.

We investigated the changes in signalling interactions to decipher how oncogenic cooperation [*Ras^V12^,scrib^-^*] promotes tumour growth. Herein, we show that the *Ras^V12^,scrib^-^* tumours grow aggressively by upregulating a previously unidentified molecular network comprising the Wingless (Wg, the *Drosophila* homolog of mammalian Wnt4), the effector caspase Dronc (*Drosophila* homolog of mammalian Caspase 9), JNK and Yki (*Drosophila* homolog of Hippo effectors YAP/TAZ in mammals). The upregulation of these signals promotes growth of *Ras^V12^,scrib^-^*tumours, and depletion of these signals reverses these phenotypes. These signals are well known for their roles in patterning and growth control (17–21), and intercellular interactions like cell competition (22–28). We show that during oncogenic cooperation these signals act together in a tumour cell-specific signalling network where Wg acts upstream of Dronc, and regulates JNK and Yki. JNK activity changes from pro-apoptosis to pro-proliferation due to a paradoxical signalling switch that ultimately promotes Yki activity. We demonstrate that in *Ras^V12^,scrib^-^*tumour cells, Yki, in turn, upregulates JNK activity causing robust activation of both JNK and Yki. This bidirectional regulatory interaction results in the formation of a self-reinforcing JNK-Yki positive feedback signal-amplification loop downstream of Wg and Dronc. We show that this molecular network is sufficient to induce tumourigenesis in other contexts of oncogenic cooperation (such as, *en*>*Yki, scrib^-^* or *Ras^V12^//scrib^-^*[interclonal tumour model (10)]). Taken together, our data strongly support the functional significance of molecular networks rather than individual signals in cancer cells, and provide novel insights into the molecular pathways and paradoxical signals that play key role in tumour growth and progression.

## 2. Materials and Methods

### 2.1 Fly stocks

The studies described in here are conducted on Drosophila melanogaster, the common fruit fly. All flies were reared in standard cornmeal-agar-molasses medium containing Tegosept and propionic acid. All fly stocks used in this study are described in Flybase (http://flybase.bio.indiana.edu). The following stocks were used in this study: Canton-S, *FRT82B*, *FRT82B scrib^j7b3^*(DGRC#111422), *scrib^dt6^*, *FRT82B scrib^2^, FRT82B scrib^2^ TubGAL-80, diap1-lacZ*, *dronc^1.7kb^-lacZ*, *ex^679^-lacZ, wg-lacZ*, *AyGAL4 UAS-GFP*, *en-GAL4*, *UAS-GFP*, *FRT82B Ubi-GFP*, *FRT82B Tub-GAL80, MS1096-Gal4, UAS-P35*, *UAS-Ras^V12^*, *UAS-sgg^S9A^, UAS-Bsk^DN^, UAS-dronc^RNAi^*, *UAS-Yki^N+CRNAi^*, *UAS-proDronc, UAS-jun^aspv^, UAS-Arm^S10^* and *UAS-Yki*.

### 2.2 Generation of somatic clones

To make MARCM clones, we crossed *eyFLP* or *UbxFLP*; *AyGAL4 UASGFP; FRT82B Tub-GAL80* virgins to males of appropriate genotypes. Heat shock mediated ‘Flp-out’ clones were generated by giving a 7min heat shock at 37°C to second instar larvae generated by crossing *AyGAL4 UASGFP* flies with *UAS Yki.* “Flp-out” clones were also generated by crossing (a) *yw hsFLP; enGAL4 ex^697^-lacZ; FRT82B UbiGFP* flies with *UAS Yki; FRT82B scrib^2^* flies (for *en>Yki; scrib^-^* clones Fig. 5). Interclonal oncogenic cooperation studies were done by crossing *yw eyFLP*; *AyGAL4 UASGFP; FRT82B Tub-GAL80 scrib^2^* virgins to *yw hsFLP; +; UASRas^V12^ FRT82B* males. All experiments were performed at 25°C unless otherwise specified.

### 2.3 Immunohistochemistry

Immunohistochemistry was done following our published protocol (29). Briefly, imaginal discs from wandering 3rd instar larvae were dissected in 1XPBS (phosphate buffered saline), fixed in 4% paraformaldehyde (PFA) for 20 minutes (min) at room temperature (RT), washed 3X10min each in 1XPBST (1XPBS + 0.2% Triton X 100), and blocked in PBSTN (PBST+ 2% Normal donkey serum) for 1h before incubation with primary antibody at 4°C overnight. The samples were then washed 3X10min each in 1XPBST, incubated in appropriate secondary antibody for two hours at RT, washed in 1XPBST for 20min and mounted in Vectashield (Vector Laboratories Inc, Burlingame, CA, USA).

The following primary antibodies were used: rabbit anti-cleaved Caspase 3 (1:250) and rabbit anti-pJNK (1:250) from Cell-Signalling Technology, Danvers, MA, USA, mouse anti-Wingless (1:100), mouse or rabbit anti-beta-galactosidase (1:100), rat anti-ELAV (1:300), and rat anti-E-cadherin (1:100) from DSHB, Iowa, IA, USA, rabbit anti-Myc (1:100) from Santa Cruz Biotechnology Inc., Dallas, TX, USA, guinea pig anti-Dronc (1:400, from Dr. H. D. Ryoo), mouse anti-DIAP1 (1:250, from Dr. B. Hayes), rabbit anti-Yorkie (1:400, from Dr. K. Irvine). The secondary antibodies were donkey Fab fragments conjugated to Cy3 or Cy5 against rabbit, mouse or guinea pig hosts (Jackson ImmunoResearch Labs, West Grove, PA, USA). We used Olympus Fluoview 1000 confocal microscope to generate projections of *z*-stacks of confocal images that were edited using Adobe Photoshop CS6.

### 2.4 Statistical analyses

Statistical analyses were performed using Microsoft Excel 2013. The magnetic lasso tool was used for clone size comparison (Fig. 1, Wild type n=11, rest n=25). Mean pixel values of clone area were obtained using the Histogram function in Photoshop CS6, and analysed using a two-tailed Student’s *t-test* assuming statistical significance at p<0.05. Intensity plots (Fig. 1, S1), were made using the plot profile function in ImageJ to find changes in pixel intensity. Dot plots were generated by calculating average signal intensity both inside and outside the clone in the Dronc or Wg channels (Fig. 1, S1) (n=5). The average wild-type values were used to find the normalization factor to determine and calculate differences in intensity levels between wild-type, *scrib^-^*, and *Ras^V12^; scrib^-^* clones. All other graphs were plotted using GraphPad Prism8.0

**Fig. 1.**
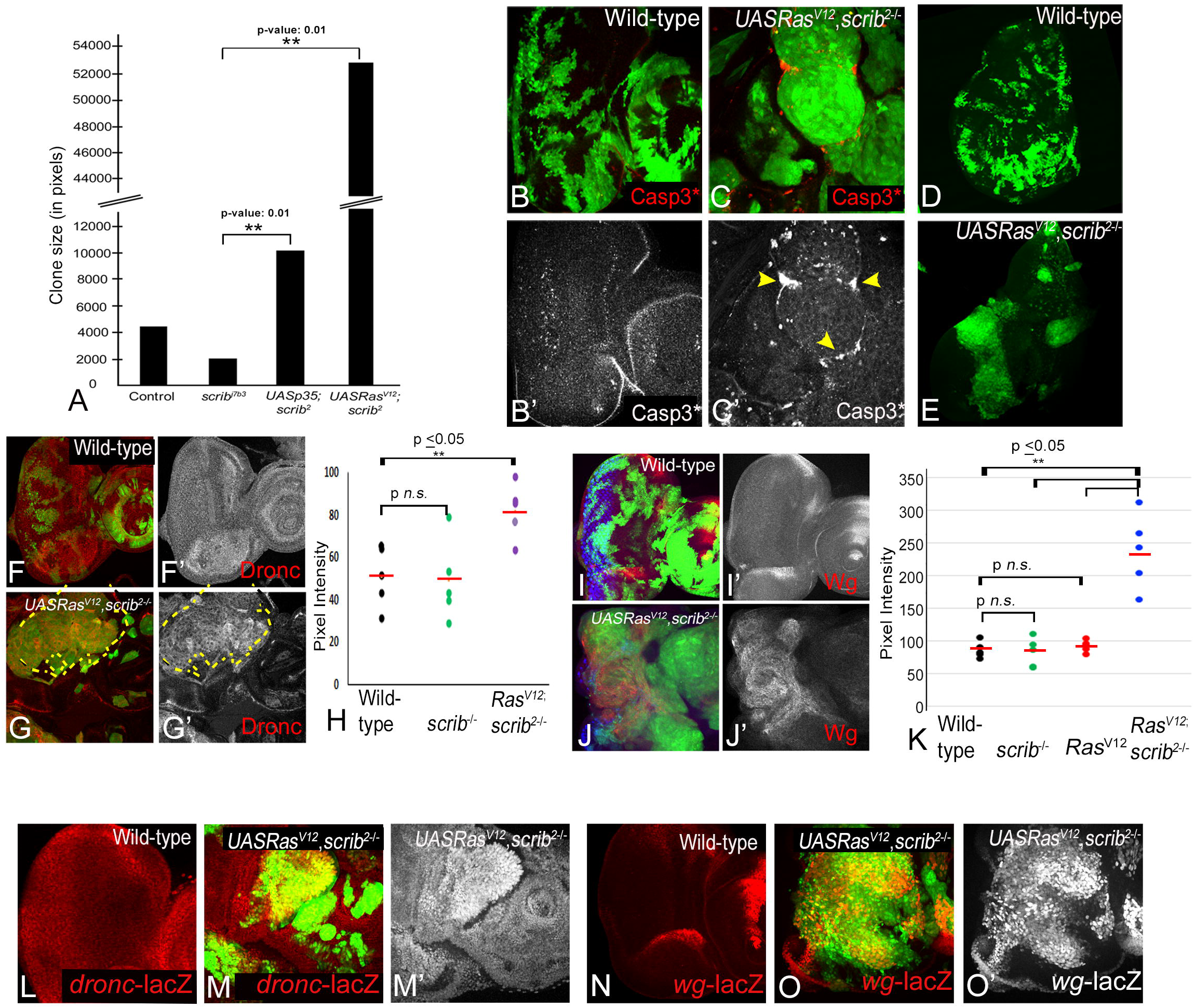
*Ras^V12^ scrib^-^* cells grow robustly and induce Dronc and Wg. (A) Graph shows quantification of clone size to comparing the growth of wildtype clones to *scrib^-^*, *p35; scrib^-^,* and *Ras^V12^ scrib^-^* clones. Student *t-test* shows significant differences (n=25, **P-value= ≤0.05). Expression of Cleaved Caspase 3 [Casp3*] antibody (red, grey) in MARCM clones (GFP, green) from third-instar eye-antennal imaginal discs (B) Wild-type, (C) *Ras^V12^ scrib^-^*, and wing discs from (D) Wild-type, (E) *Ras^V12^ scrib^-^*. Yellow arrowheads mark dying cells in C’. (F-H) Panels show Dronc expression (red, grey) in MARCM clones (green) induced in eye-antennal imaginal discs from (F-F’) wild-type, and (G, G’) *Ras^V12^ scrib^-^*. (H) Dot blot comparing Dronc expression (signal intensity) in MARCM clones in the indicated genotypes (n=5, **P-value= ≤0.05). (I-K) Expression and quantification of Wingless expression in eye discs from (I, I’) wild-type, and (J, J’) *Ras^V12^ scrib^-^* MARCM clones (green). (K) Dot blot shows change in Wg expression (signal intensity) in the indicated genotypes (n=5, **P-value= ≤0.05). The horizontal red line marks the cumulative mean expression levels in panels H and K. (L-M’) *dronc^1.7kb^-lacZ* reporter expression (red, grey) in (L) wild-type and (M, M’) *Ras^V12^ scrib^-^* MARCM clones (green). (M-O’) *wg-lacZ* reporter expression (red, grey) in (L) wild-type and (M, M’) *Ras^V12^ scrib^-^* MARCM clones (green). The magnification and orientation of discs is identical in all panels.

### 2.5 qRT-PCR

RNA was extracted from eye imaginal discs from wandering third instar larvae. At least 40-50 discs were collected per genotype in TRIzol. The ZYMO RNA Clean and Concentrator Kit was used for RNA extraction. To prepare cDNA, 200ng/ul RNA/sample was reverse transcribed using the cDNA synthesis kit (GE, Cytiva). qRT-PCRs were done in triplicate per sample using the iQ SYBER Green Supermix (BIORAD) on an iCycler iQ™ Real-Time PCR Detection System. Three biological and three technical repeats were done for each experiment, and data analyzed using the ddCt method.

Primers (IDT) used were:

***wg*:** *Forward primer: 5’* CGT CAG GGA CGC AAG CAT A −3’

*Reverse Primer: 5’* ATT GTG CGG GTT CAG TTG G −3’

***dronc*:** *Forward primer: 5’* CGA TGG ATC TGT GGT CGA TAT G −3’

*Reverse Primer: 5’* GGC TTC GCT CGT CTT CTT TA −3’

***Gapdh****: Forward primer: 5’* TAA ATT CGA CTC GAC TCA CGG T −3’

*Reverse Primer: 5’* CTC CAC CAC ATA CTC GGC TC −3’

## Results

### 3.1. Ras^V12^,scrib^-^ cells grow robustly and induce JNK, Yki, Dronc and Wg

To investigate the changes in intercellular interactions that promote tumour growth, we made *ey-FLP* and *Ubx-FLP* induced MARCM clones [GFP positive] in the eye (Fig. 1A-C) and wing discs (Fig. 1D, E) respectively and monitored clone size to assess tumour growth (Fig.1A). Compared to wild-type, *scrib^-^* clones grew poorly (Fig. 1A)(30). Given that *scrib^-^* clones are competed out due to cell competition mediated apoptosis (Fig. S1A), we blocked apoptosis by overexpressing UASp35 (Fig. S1B), a pan-Caspase inhibitor(31). The inhibition of apoptosis improved the growth of *scrib-* clones (*scrib^-^,P35,* Fig.1A, P-value: 0.01), however the discs remain monolayered and clones did not form tumours (Fig. S1B). In contrast, coexpression of oncogenic Ras in *scrib^-^* cells (*Ras^V12^,scrib^-^*) resulted in robust aggressive and invasive tumours(8) that grew several fold compared wild-type, *scrib^-^*, or *scrib^-^*,*p35* clones (Fig.1A, P-value: 0.01). These data suggest that a paradoxical switch changes the ability of *scrib^-^* cells, and inter-cellular interactions may be a critical part of these observed effects. Next, to understand the nature of intercellular interactions that may underlie the differences in clone sizes, we tested cell death by using an antibody against activated Caspase 3 [Casp3*]. In comparison to wild-type clones (Fig. 1B-B’), in *Ras^V12^,scrib^-^*clones a bulk of cell death was induced in the wild-type cells surrounding the clone (Fig. 1C, C’). Similarly in the wing discs, compared to wild-type (Fig.1D) *Ras^V12^,scrib^-^*clones grew to large invasive tumours (Fig. 1E). Taken together, these data suggested that the *Ras^V12^,scrib^-^* clones grow robustly, therefore, we tested if JNK, Yki, Dronc and Wg, previously linked to cell survival and proliferation during cell competition, were affected in the *Ras^V12^ scrib^-^* tumours.

JNK and Yki are known to interact in a context-dependent manner (12–14,32), and increased JNK or Yki activity has been linked to tumour growth (12,33–35). In contrast, elimination of *scrib^-^*cells by cell competition involved JNK-dependent suppression of Yki activity (12,30,36–38). To test Yki activity, we employed the Yki-reporter *diap1-lacZ,* which is expressed ubiquitously in wild-type (Fig. S1C) discs. Increased Yki activity due to Yki overexpression in ‘flp-out’ (*AyGAL4>Yki*) clones showed robust induction of *diap1-lacZ* (Fig. S1D). In the *Ras^V12^,scrib^-^* clones *diap1-lacZ* was strongly induced (Fig. S1E,E’). Interestingly, *diap1-lacZ* was also induced in peri-tumoural cells (non-cell autonomously in wild-type cells) abutting the *Ras^V12^,scrib^-^* tumours (Fig. S1E’). The pixel intensity from controls and experimental groups was quantified, and showed significant increase in cells Yki overexpression and in *Ras^V12^,scrib^-^* clones (Fig. S1F). Compared to wild-type (Fig. S1G, G’), pJNK was robustly induced in the *Ras^V12^,scrib^-^* clones (Fig. S1H,H’, quantified in I). Thus, consistent with previous data, JNK and Yki activities were simultaneously upregulated in the *Ras^V12^,scrib^-^* tumours. We then investigated the signals that converge onto Yki and JNK to promote growth of *Ras^V12^,scrib^-^* tumours.

Previous studies have shown that Dronc (the initiator caspase that induces apoptosis through activation of Caspase 3) and Wg (the secreted ligand of the Wingless/Wnt pathway that acts as a mitogen) play critical roles in intercellular interactions like cell competition and compensatory proliferation (27,39–41). Yki also transcriptionally regulates *dronc* and *wg* (42,43). We tested levels of active Dronc using an antibody against the activated Dronc^CA^ form (41), and Wg in the *Ras^V12^,scrib^-^* clones (Fig. 1). Dronc is expressed ubiquitously in wild-type imaginal discs (Fig. 1F, F’, quantified in Fig. 1H), and its levels remained unaltered in *scrib^-^* cells in wild-type background (Fig. 1H) suggesting that wild-type Dronc levels were sufficient for elimination of *scrib^-^* cells in the presence of elevated pJNK levels. In comparison, Dronc was strongly upregulated in *Ras^V12^,scrib^-^*cells (Fig. 1G, G’). Quantification of signal intensities showed that Dronc levels were significantly upregulated (1.7 fold) in *Ras^V12^,scrib^-^* cells compared to either wild-type or *scrib^-^* cells (Fig. 1H). Taken together, these data suggested that Dronc plays a non-apoptotic role and promotes *Ras^V12^,scrib^-^* tumour growth. In wild-type eye discs, Wg is expressed at the lateral margins anterior to the morphogenetic furrow (Fig. 1I, I’, quantified in K) (44). This pattern of Wg expression was unaltered in *scrib^-^* cells generated in wild-type background (Fig. 1K). In *Ras^V12^,scrib^-^* cells, Wg was robustly induced (Fig. 1J, J’). When compared to wild-type or *scrib^-^* cells, mean Wg expression levels showed a 2.7 fold increase in *Ras^V12^,scrib^-^* cells (Fig. 1K) suggesting that the steep upregulation of Wg levels in *Ras^V12^,scrib*^-^ may itself promote super-competition by promoting proliferation through its mitogenic functions.

Next, we checked if *dronc* and *wg* transcription was affected in *Ras^V12^,scrib^-^* cells using the *dronc^1.7kb^-lacZ* (Fig. 1L-M’) and *wg-lacZ* (Fig. 1N-O’) reporters. We found that compared to the wild-type (Fig. 1L), *dronc^1.7kb^-lacZ* expression was robustly induced in the *Ras^V12^,scrib^-^* clones (Fig. 1M,M’). Similarly, the normal pattern of *wg-lacZ* expression in eye discs (Fig. 1N) was disrupted due to ectopic induction of *wg-lacZ* in *Ras^V12^,scrib^-^* clones (Fig. 1O,O’). We confirmed the increased expression of *dronc* and *wg* in *Ras^V12^,scrib^-^* clones using qRT-PCR (Fig. S2). The upregulation of *dronc* and *wg* in the tumour cells suggests that these genes promote tumour growth possibly by co-opting compensatory mechanisms where *dronc* plays a non-apoptotic paradoxical role and *wg* plays a mitogenic role. Taken together, these data suggest that four signals associated with intercellular interactions (Yki, JNK, Wg, Caspases) were upregulated in the aggressively growing *Ras^V12^,scrib^-^* tumours.

### 3.2. Ras^V12^,scrib^-^ tumours require JNK, Yki, Dronc and Wg for growth

To understand if some or all of these signals played a key role in the aggressive growth of *Ras^V12^,scrib^-^* tumours, we tested their requirement by downregulating these signals in *Ras^V12^,scrib*^-^ clones (Fig.2) using transgenes that downregulated Dronc [*Dronc^RNAi^*](42), JNK [*Bsk^DN^*](30), Wg [Sgg^S9A^](45) and Yki [*Yki^N+CRNAi^*](46). A comparison of clone sizes revealed that individual downregulation of the four signalling pathways significantly decreased the growth of the *Ras^V12^,scrib^-^*cells (Fig. 2A, B-E). These data suggested a requirement of the four signals for the aggressive growth of *Ras^V12^,scrib^-^* tumours. Therefore, we next tested if loss of cellular fitness or interactions amongst these signals was the underlying cause for the change in tumour size.

**Fig. 2.**
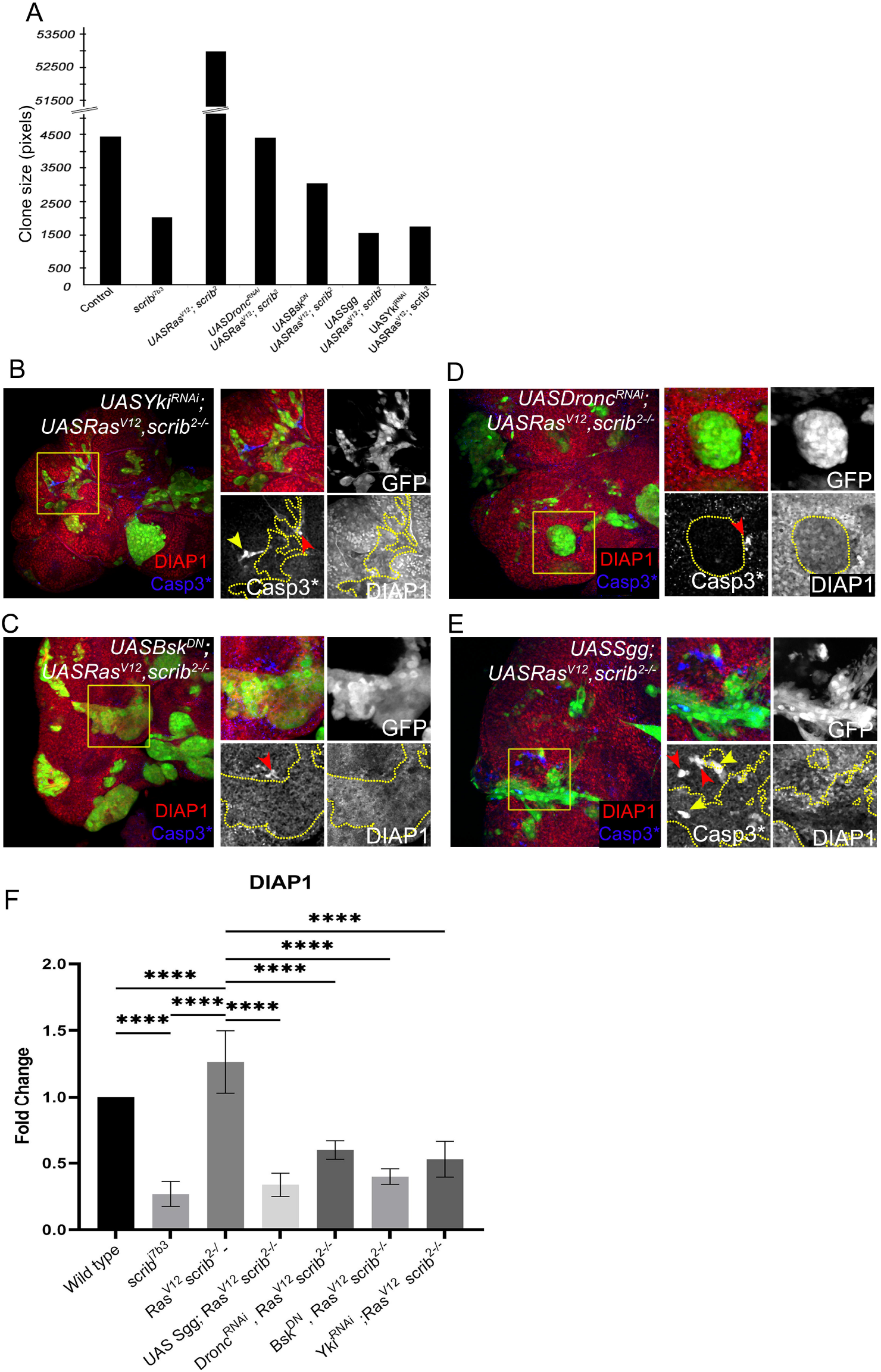
*Ras^V12^ scrib^-^* tumours require Dronc, Wg, JNK and Yki for growth. (A) Bar graph shows a quantification of clone size of the following genotypes: Wild-type, *scrib^-^*, *Ras^V12^ scrib^-^, UASDronc^RNAi^ UASRas^V12^ scrib^-^*, UAS*Bsk^DN^ UASRas^V12^ scrib^-^*, *UASSgg^S9A^ UASRas^V12^ scrib^-^*, *UASYki^N+CRNAi^ UASRas^V12^ scrib^-^* (n=5, **P-value= ≤0.05) (B-E) Panels show DIAP1 (red) and Casp3* (blue) expression in eye imaginal discs containing MARCM clones (green) of the following genotype: (B) *UASYki^N+CRNAi^ UASRas^V12^ scrib^-^*, (C) *UASBsk*^DN^ *UASRas^V12^ scrib^-^*, (D) *UASDronc^RNAi^ UASRas^V12^ scrib*, and (E) *UASSgg^S9A^ UASRas^V12^ scrib^-^.* Clone areas highlighted by yellow boxes are further magnified in the image to the right and split channel images in grescale are presented for each genotype. Clone outlines are marked by the yellow dotted line, red arrowheads highlight Casp3* positive apoptotic cells outside the clones, whereas yellow arrowheads mark Casp3* positive apoptotic cells inside the clones. (F) Graph presents the fold change in DIAP1 levels comparing wild-type and *UASRas^V12^ scrib^-^* to all tested genotypes. Error bars show SEM (standard error of means) and significant differences quantified using ordinary one-way ANOVA (****P-value=≤0.001). The magnification and orientation of discs is identical in all panels.

### 3.3 Downregulation of JNK, Yki, Dronc and Wg impairs cellular fitness

We hypothesized that the reduction in clone size was due to altered fitness within *Ras^V12^,scrib^-^*cells. First, we tested changes in DIAP1 levels (Fig. 2B-E red, grey) in conditions where at least one of these four signals were downregulated (Fig. 2B-F). DIAP1 inhibits caspase activation, and protects cells from caspase-mediated apoptosis(47). DIAP1 was ubiquitously expressed in wild-type cells (Fig. S3A), downregulated in *scrib^-^* cells (Fig. S3B, yellow arrowheads), and robustly induced in *Ras^V12^,scrib^-^* clones (Fig. S3C). However, when either Yki (Fig.2B), JNK (Fig. 2C), Dronc (Fig.2D), or Wg (Fig. 2E) were downregulated in *Ras^V12^,scrib^-^*clones (Fig. 2C-E, yellow outline in grey channels), DIAP1 was downregulated (quantified in Fig. 2F). This suggests that the *Ras^V12^,scrib*^-^ tumour cells had decreased survival, and were susceptible to elimination by apoptosis (Fig. 2B-E, blue, grey) when the tumour promoting signals were downregulated. Indeed, downregulation of Yki (Fig. 2B) or Wg (Fig. 2E) resulted in apoptosis both inside (Fig. 2B, E yellow arrowheads) and outside (Fig.2B, E, red arrowhead) the *Ras^V12^,scrib^-^*clones suggesting that Wg and Yki protect *Ras^V12^,scrib^-^*tumours during competitive interactions. Downregulation of JNK pathway (Fig. 2C) or Dronc (Fig. 2D) resulted in apoptosis at the clone boundary (Fig. 2C-D, red arrowhead) but not within the clones likely because the cell death regulators (JNK and Dronc) were suppressed in the clones. Taken together, these data suggested that downregulation of the tumour promoting signals impaired cellular fitness in *Ras^V12^,scrib^-^* cells.

These observations were further supported in control experiments, where we made MARCM clones in which we depleted each component of the molecular network in otherwise wildtype cells (Fig. S4). Downregulation of Wg signalling (confirmed with anti-Wg antibodies, Fig. S4B red, A”’ grey) caused cell death (Fig. S4A blue, A”’ grey) and downregulation of DIAP1 (Fig. S4A red, A”’ grey). However, the levels of pJNK (Fig. S4B blue, A”’ grey) remained unaltered. Depletion of Dronc caused a block in cell death (Fig. S4C blue, A”’ grey) and did not significantly affect DIAP1 (Fig. S4C red, A” grey), Wg (Fig. S4D red, A” grey) or pJNK (Fig. S4C blue, A”’ grey) expression. Blocking JNK signalling (Fig. S4F blue, A”’ grey) showed similar effects as downregulation of Dronc on DIAP1 (Fig. S4E red, E” grey), Wg (Fig. S4F red, F” grey) or pJNK (Fig. S4F blue, A”’ grey). Depletion of Yki resulted in downregulation of DIAP1 (Fig. S4G red, G” grey) and a strong non-cell-autonomous induction of cell death (Fig. S4G blue, G”’ grey). No significant effects on Wg (Fig. S4H red, H’ grey) or pJNK (Fig. S4H blue, H’ grey) were seen. Thus, Wg and Yki emerged as key signalling proteins amongst these four pathways.

### 3.4 Wg, Dronc, JNK and Yki form a molecular network in Ras^V12^,scrib^-^ tumours

Next, we tested if Wg, Dronc, JNK and Yki regulated one another to promote the aggressive growth of *Ras^V12^,scrib^-^* tumours (Fig. 3). Downregulation of Wg signalling in *Ras^V12^,scrib^-^* cells (Fig. 3A) resulted in small tumours and reduction in Wg expression (Fig. 3A red, A” grey). Interestingly, depleting Dronc (Fig. 3B), JNK (Fig. 3C), or Yki (Fig. 3D) caused a significant decrease in clone size, but Wg was strongly induced in these clones (Fig., B”-D”, quantified in Fig. 3E). Thus, Wg acts upstream of Dronc, JNK and Yki in the network of signals that promote *Ras^V12^,scrib^-^*tumour growth.

**Fig. 3.**
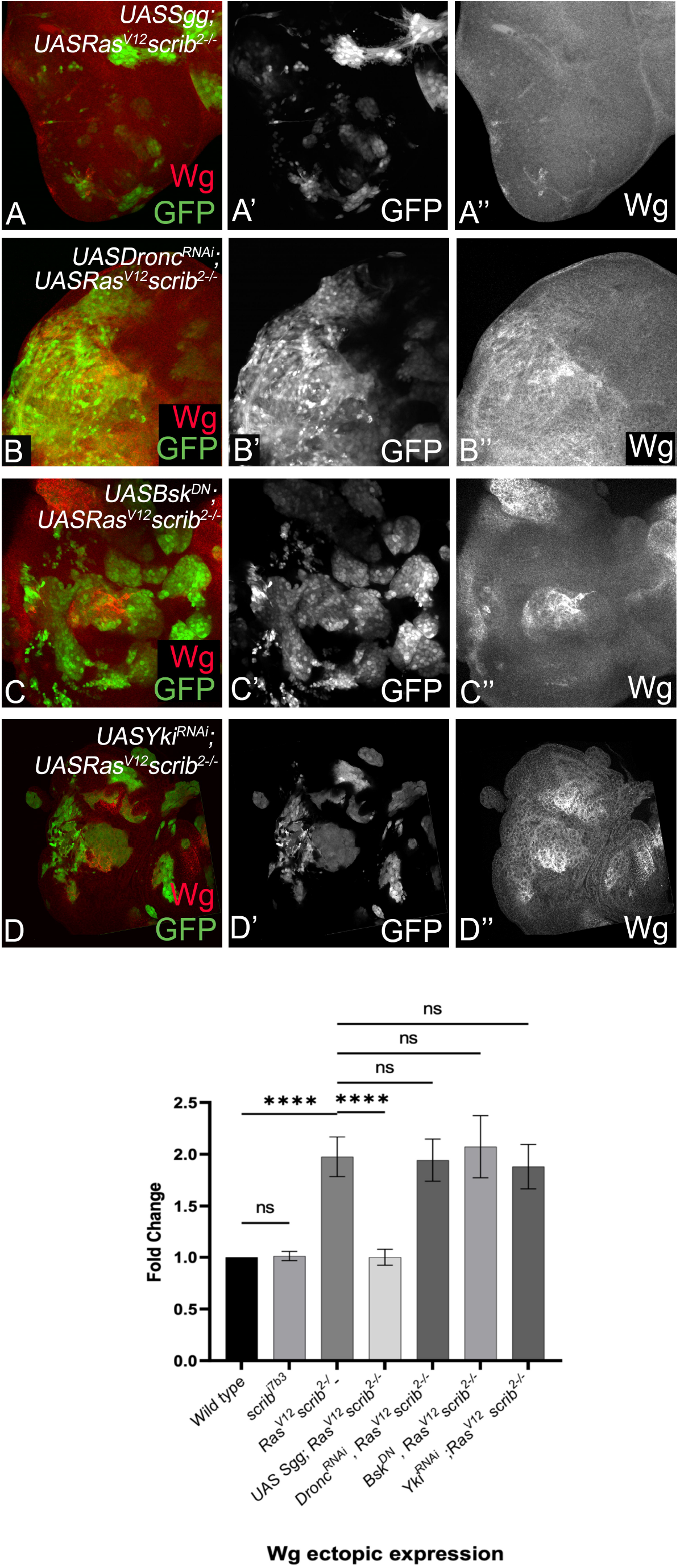
Wg acts upstream of the molecular network. Panels show Wg expression (red A-D, grey A”-D”) in MARCM clones (green in A-D, grey in A’-D’) of the following genotype: (A) *UASSgg^S9A^ UASRas^V12^ scrib^-^*, (B) *UASDronc^RNAi^ UASRas^V12^ scrib^-^*, (C) *UASBsk^DN^ UASRas^V12^ scrib^-^*, and (D) *UASYki^N+CRNAi^ UASRas^V12^ scrib^-^.* Graph presents the fold change in Wg levels comparing wild-type and *UASRas^V12^ scrib^-^* to all tested genotypes. Error bars show SEM (standard error of means) and significant differences quantified using unpaired t-test (****P-value=≤0.001). The magnification and orientation of discs is identical in all panels.

To test if these signals interact with each other, we checked effects of loss of Yki (*Yki^RNAi^*;*Ras^V12^,scrib^-^*, Fig. 4A-D) on Dronc (Fig. 4A), and pJNK expression (Fig. 4B). Depletion of Yki (confirmed in Fig. 4C red, C’ grey) did not affect induction of Dronc (Fig. 4A blue, A’ grey), however pJNK expression was downregulated in these clones (Fig. 4C blue, C’ grey). We quantified the mean grey values and plotted the fold change comparing Wild-type to *Ras^V12^,scrib^-^* and *Yki^RNAi^; Ras^V12^,scrib^-^*, and the graphs show significant downregulation of Yki and pJNK, whereas Dronc accumulation was not affected (Fig. 4D). Next, we downregulated JNK signalling (Fig. 4E-G) by overexpression of a dominant negative form of Basket (Bsk) in *Ras^V12^,scrib^-^* (*Bsk^DN^;Ras^V12^,scrib^-^*) which results in downregulation of pJNK (Fig. 4E blue, E’ grey), but Dronc expression remained upregulated in these clones (Fig. 4E red, E’ grey). Interestingly, Yki expression was also depleted in *Bsk^DN^;Ras^V12^,scrib^-^* cells (Fig. 4F red, F’ grey). The fold change in expression of Dronc, Yki and pJNK confirms these effects (Fig. 4G). Interestingly, downregulation of Dronc (*Dronc^RNAi^;Ras^V12^,scrib^-^*, Fig. 4H-J) in *Ras^V12^,scrib^-^* cells (Fig. 4H red, H’ grey) showed downregulation of JNK (Fig. 4H blue, H’ grey) and Yki (Fig. 4I red, I’ grey). Quantification shows that compared to wild-type or *Ras^V12^,scrib^-^*significant downregulation of Yki and JNK is seen in *Dronc^RNAi^;Ras^V12^,scrib^-^*clones (Fig. 4J). Downregulation of Wg (Fig. 4K-M) in *Ras^V12^,scrib^-^* cells (Sgg^S9A^;*Ras^V12^,scrib^-^*) showed downregulation of Dronc (Fig. 4K red, K’ grey), pJNK (Fig. 4K blue, K’ grey) and Yki (Fig. 4L red, L’ grey). Quantification of effects of downregulation of Wg revealed significant downregulation Dronc, Yki and pJNK (Fig. 4M). These data show that Dronc levels remain upregulated and comparable to *Ras^V12^,scrib^-^* cells in combinations where either Yki (Fig. 4A-D) or JNK (Fig. 4E-G) are downregulated, suggesting that Dronc acts upstream of Yki and JNK in this signalling network. Furthermore, JNK and Yki regulate each other suggesting that they may act in a feedforward loop downstream of Dronc. Overall, our data suggest that the growth of the *Ras^V12^,scrib^-^* clones depended on a network involving Wg dependent activation of Dronc, which controls a JNK-Yki mediated signal amplification loop that sustains high levels of JNK and Yki activities.

**Fig. 4.**
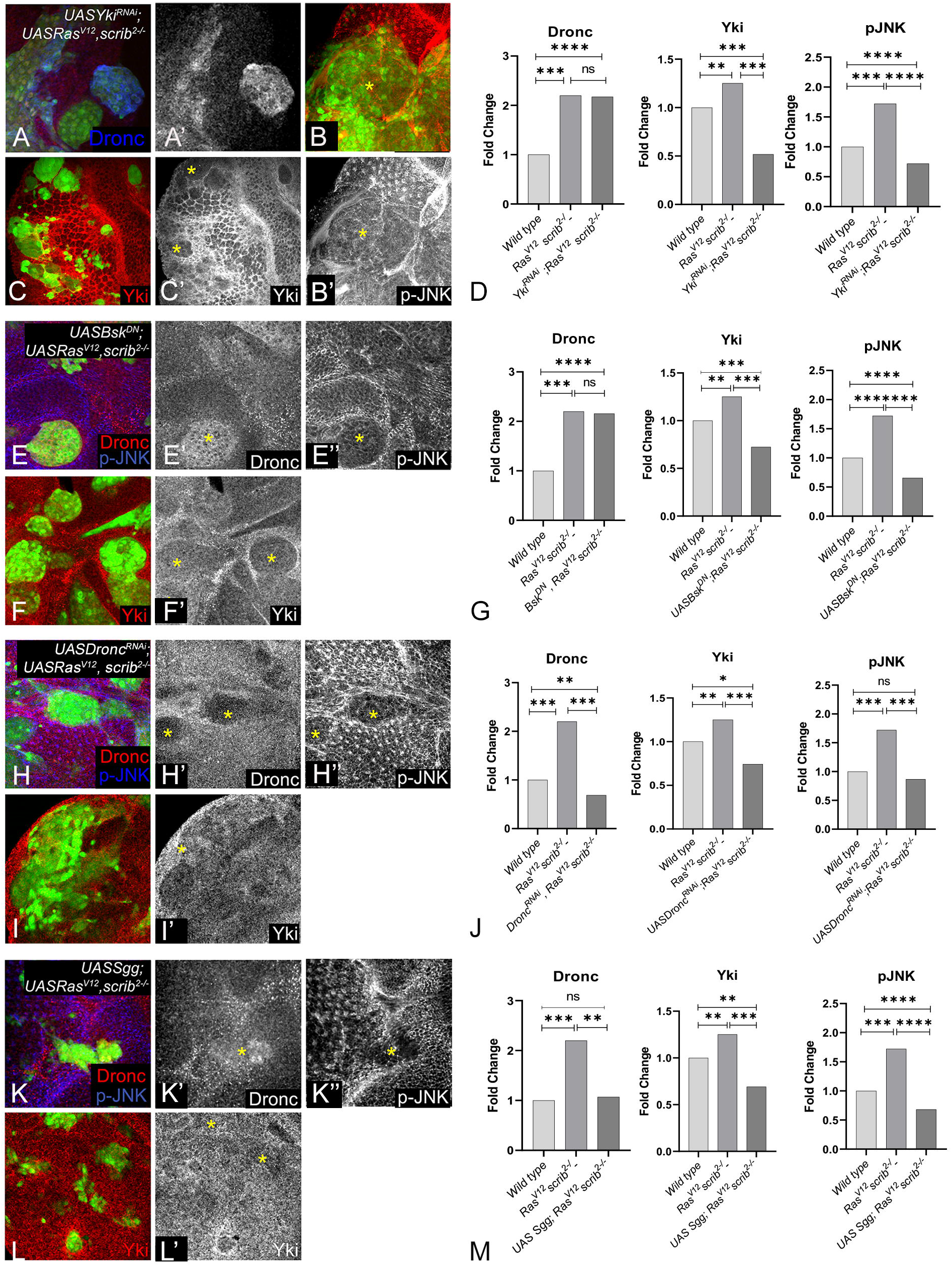
Wg, Dronc, JNK and Yki form a molecular network in *Ras^V12^ scrib^-^* clones. Panels show Dronc, Yki and p-JNK expression in MARCM clones (green) when the molecular network is perturbed. (A-D) Panels show *UASYki^N+CRNAi^ UASRas^V12^ scrib^-^* clones stained for Dronc (A blue, A’ grey), pJNK (B red, B’ red) and Yki (C red, C’ grey), quantified in D. (E-G) *UASBsk^DN^ UASRas^V12^ scrib^-^* clones showing expression of Dronc (E red, E’ grey), pJNK (E blue, E” grey), and Yki (D red, F’ grey), quantified in G. (H-J) Confocal images of *UASDronc^RNAi^ UASRas^V12^ scrib^-^* clones showing expression of Dronc (H red, H’ grey), pJNK (H blue, H” grey), and Yki (I red, I’ grey) are presented. The quantification of fold-change is shown in J. (K-M) Panels show confocal scans of *UASSgg^S9A^ UASRas^V12^ scrib^-^* clones stained for Dronc (K red, K’ grey), pJNK (K blue, K” grey), and Yki (L red, L’ grey), quantified in M. The quantifications in D, G, J, M show fold change comparison between indicated genotypes, error bars show SEM (standard error of means) and significant differences quantified using unpaired t-test (****P-value=≤0.001). The magnification and orientation of discs is identical in all panels.

### 3.5 Cooperative interactions stimulate the molecular network and tumour growth

The preceding data led us to ask, if oncogene activation, loss of polarity (scrib^-^) or cooperative interactions were important in stimulating the molecular network and tumour growth. Loss of polarity (*scrib*) alone is insufficient to induce aggressive growth in somatic clones (Fig. 1, S1) (8,9,12,30), therefore, we tested the importance of cooperative interactions on induction of the Wg-Dronc-JNK-Yki network and tumour growth. We tested two scenarios where loss of polarity could synergize with oncogene activation.

First, we generated *scrib^-^* clones [GFP negative, generated by hs-FLP] in wing discs where Yki was overexpressed in the posterior compartment [*en>Yki; scrib^-^*] (Fig. 5). We chose *en>Yki; scrib^-^* combination because loss of *scrib* alone in eye or wing discs were susceptible to elimination by cell competition. In the eye discs (as in the wing discs) *scrib* upregulated JNK (Fig. S5A-B’), downregulated Yki (Fig. S5C-D’) and had little effect on Dronc (Fig. S5E) or Wg (Fig. S5F) expression. Thus, loss of *scrib* had clear and distinguishable effects from Yki overexpression (Fig. 6A) on the four signals and provided an ideal opportunity to test altered signalling during oncogenic cooperation. We found that *scrib^-^* cells grew to significantly larger sizes in the posterior (P) compartment where Yki was overexpressed (Fig. 5A-E), and showed reduced E-cadherin expression (Fig. 5A, red, grey, red asterisk). In addition to the increased Yki activity (monitored by *ex-lacZ* expression) in the posterior compartment expected in response to Yki overexpression (Fig. 5C), we saw that Yki activity was very strongly induced within and around the *scrib^-^* cells (Fig. 5C, outlined by yellow line). We called this dramatic increase in Yki activity ‘super-induction’. In these clones, MMP1 (Fig. 5B red, grey), pJNK (Fig. 5D red, grey), and Wg (Fig. 5E red, grey) were induced in and around the *scrib^-^* clones, potentially leading to the establishment of the JNK-Yki loop and invasiveness by induction of MMP1. The Yki, pJNK and Wg signals spread to several cells outside the *scrib^-^* clones with Yki activity propagating the farthest (Fig. 5C). In contrast, in the anterior (A) compartment *scrib^-^* clones (Fig. 5B-E blue outline) behaved similar to clones in wild-type background (Fig. S1A), and were eliminated by cell competition. Comparison of the P and A compartment-specific clones suggested that JNK-Yki signal amplification loop was activated specifically in the tumour cells in the P compartment where cooperative interactions occurred between Yki-expressing and *scrib^-^* cells.

**Fig. 5.**
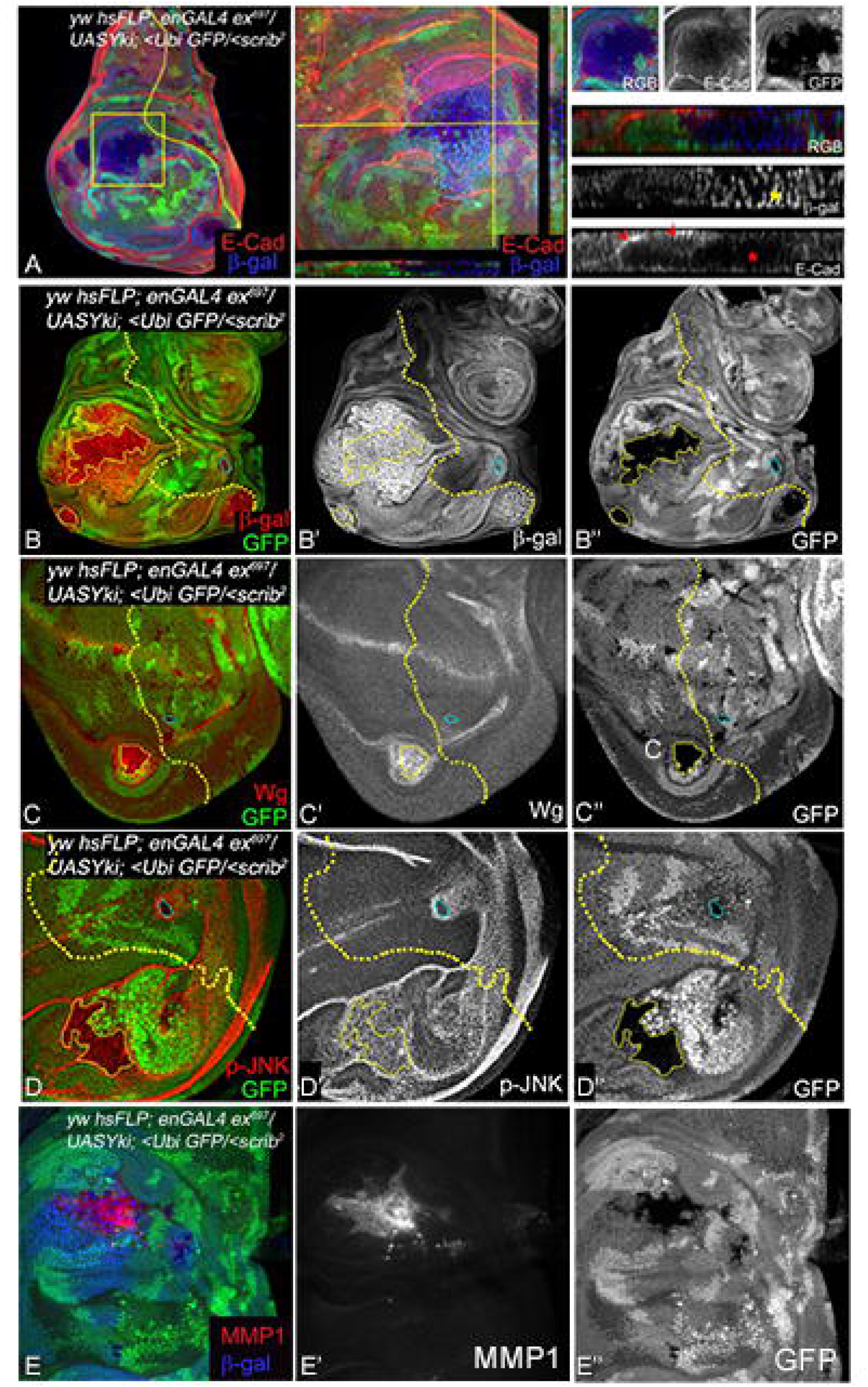
Cooperative interactions induce the Wg-Dronc-JNK-Yki network to stimulate tumour growth. (A) Wing imaginal discs showing *en>Yki, scrib^-^* clones (GFP-negative) stained for *ex-lacZ* (using antibodies to β-gal, A, B red) and E-cadherin (blue). The area highlighted by the yellow box is shown at higher magnification in the panels to the right. Cropped image of the clone is shown next to depict changes in polarity and growth. Red asterisks point towards the loss of E-cadherin in the XZ section (along the plain marked in the middle panel), and normal E-cadherin localization is highlighted by red arrows for comparison. (B) Yki activity as assessed by β-gal (blue) expression (B red, B’ grey) in *en>Yki, scrib^-^* clones is shown. (C-E) Wing discs containing *en>Yki, scrib^-^* clones (GFP negative) were assayed for Wg (C red, C’ grey), p-JNK (D red, D’ grey) or MMP1 (E red, E’ grey) expression. The *scrib^-/-^* clones in posterior compartment (solid yellow line) and anterior compartment (blue lines) are highlighted for each marker. In all images the anterior posterior compartment boundary is marked by yellow dotted line. Images in A-B are at 20X magnification, and C-E are at 40X magnification. Posterior is to the left and dorsal up in all discs.

**Fig. 6.**
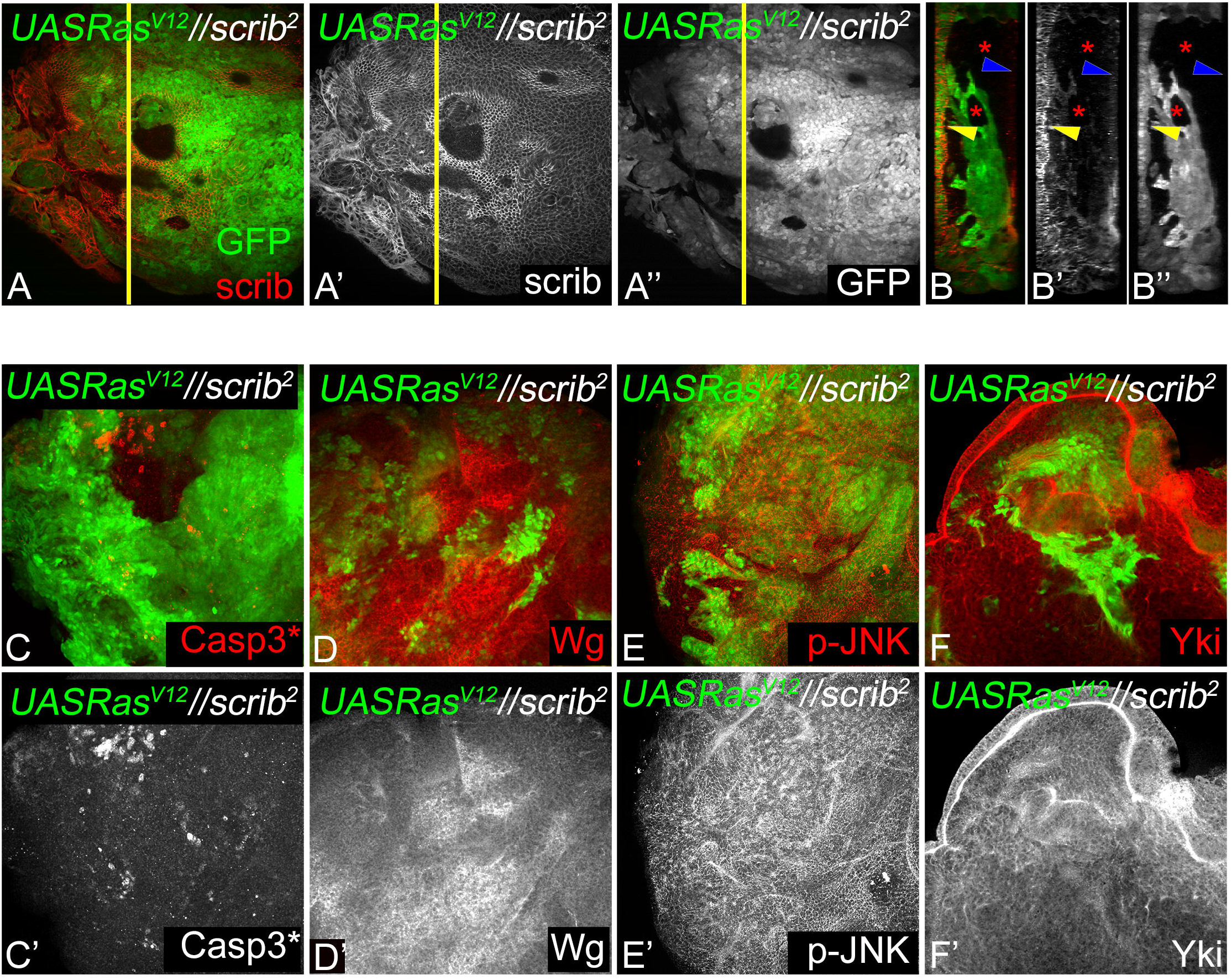
Wg-Dronc-JNK-Yki network drives tumour growth during interclonal interactions. (A) *Ras^V12^//scrib^-^* interactions in eye discs show small *scrib^-^* clones identified by loss of Scrib (red) and adjacent *Ras^V12^* clones (GFP, green). (B-B”) The YZ projection of the yellow line in A shows a multi-layered *Ras^V12^* clone (GFP). (C-F) Eye discs show expression of Casp3*(C red, C’ grey), Wg (D red, D’ grey), p-JNK (E red, E’ grey), and Yki (F red, F’ grey),) in *Ras^V12^//scrib^-^*interclonal interactions. Note upregulation of Casp3*, Wg and Yki at the boundary of *Ras^V12^* clones.

Second, we tested if the Wg-Dronc-JNK-Yki network was activated during interclonal cooperation when *Ras^V12^* and *scrib^-^* cells were adjacent to each other (*Ras^V12^//scrib^-^*)(10). Consistent with previous data, aggressive *Ras^V12^* tumours (GFP positive) were formed adjacent to smaller *scrib^-^* clones (Fig. 6A)(10). Interestingly, the Ras*^V12^* clones extruded apically and showed a multi-layered invasive phenotype (Fig. 6A-A”, Y-Z sections). The Ras clones showed robust growth despite induction of cell death at the clone boundary between Ras*^V12^* (GFP-positive) and *scrib* (GFP-negative) (Fig. 6B, B’) clones. Expression of Wg was robustly induced in the *scrib^-^* clones (Fig. 6C, C’) whereas pJNK levels appear upregulated in both *scrib^-^* and *Ras^V12^* clones (Fig. 6D, D’)(10,30). Taken together these data suggest that the robust growth of the *Ras^V12^* clones was dependent on the interclonal signalling interactions between Caspases, Yki, Wg and JNK signalling emanating from the *scrib^-^*clones. Overall, these two experimental approaches reaffirmed that the induction of the molecular network was intimately linked to tumor growth in diverse scenarios.

### 3.6 The role of apical-basal polarity in the establishment of tumour specific molecular networks

The requirement of the Wg-Dronc-Yki-JNK network in promoting tumour growth under various cooperative contexts raised the question if these four signals were sufficient to induce tumours in normal cells or loss of polarity is essential to form aggressive tumours. To test this, we first tested the effect of overexpressing Wg-Dronc-Yki-JNK in normal cells using the Gal4-UAS system in wing-imaginal discs. We used the *MS1096-Gal4* (Fig. 7A) to drive expression of transgenes overexpressing *Yki, pro-Dronc, jun^aspv^, and Arm^S10^* (MS1096>*Yki,pro-Dronc, jun^aspv^, Arm^S10^*) which would result in activation of all four signals (Fig. 7). Interestingly, co-expression of these transgenes resulted in hyperplasia of the wing pouch and hinge region (Fig. 7B-B”). We observed moderate upregulation of DIAP1 in the wing pouch region (Fig. 7B’) and increased cell death in the wing hinge region (Fig. 7B”). Both Wg (Fig. 7C, C’) and pJNK are induced in a patchy pattern in the dorsal hinge region (Fig. 7C, C”). In control experiments, overexpression of individual transgenes revealed that overexpression of *Yki* (*MS1096>Yki*) is capable of driving hyperplasia by upregulation of DIAP1, Wg and pJNK (Fig. S6A-B”). Overexpression of pro-Dronc (*MS1096>proDronc*) did not affect the growth of the disc very strongly and showed moderate upregulation of DIAP1, cell death, Wg and pJNK (Fig. S6C-D”) suggesting that pro-Dronc plays a role in promoting survival but not cell proliferation. Overexpression of *jun^aspv^*(*MS1096>jun^aspv^*) showed the most interesting effects by reducing size of the wing pouch due to strong suppression of DIAP1 and induction of cell death (Fig. S6E-F’’). Although Wg and pJNK are induced in this combination, the excessive cell death in wing pouch does not support the growth of these discs. Activation of Wg pathway by overexpression of Arm^S10^ (*MS1096>Arm^S10^*) resulted in upregulation of DIAP1, Wg and pJNK (Fig S6G-H”). In addition, mild to moderate levels of cell death were observed in the wing pouch. Taken together, these data suggest that although coactivation of these transgenes results in increased growth, the presence of normal wild-type cells with intact polarity do not allow the establishment of the network that would drive the tumour growth.

**Fig 7.**
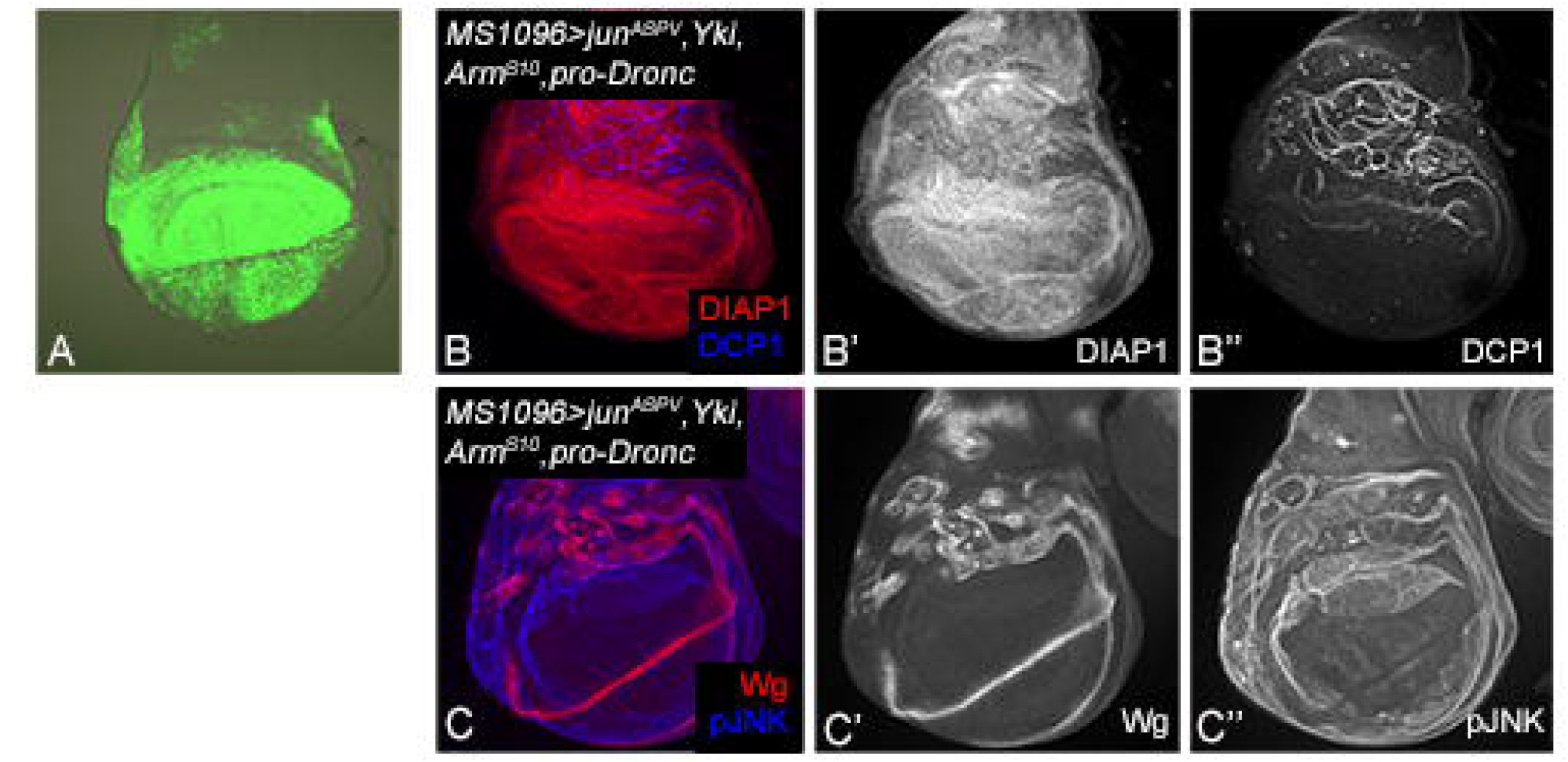
Loss of polarity is required for the Wg-Dronc-JNK-Yki network to drive tumour growth. (A) Control wing imaginal disc from *MS1096 Gal4>UASGFP* is shown. (B-C”) Panels show wing imaginal discs from *MS1096 Gal4>UASjun^aspv^, UASYki, UASproDronc, UASArm^S10^* stained for expression of DIAP1 (B red, B’ grey), DCP1 (B blue, B” grey), Wg (C red, C” grey) and pJNK (C blue, C” grey). The magnification and orientation of discs is identical in all panels.

## 4. Discussion

Oncogenic cooperation is a key mechanism for tumour development and progression. Cooperative interactions between oncogenic *Ras* and loss of *scrib^-^* (*Ras^V12^,scrib^-^*) have been elegantly modelled using *in-vivo* mosaic tumour models in *Drosophila* (8,9,12,30), and multiple mammalian cancer models (48–52). We investigated how altered signalling and cell-cell interactions promote tumourigenesis during oncogenic cooperation. Previously ectopic upregulation of a network of transcription factors e.g., Upd, JAK-STAT, AP-1, Myc, Ftz, Ets, Irbp18, Xrp1, Slow border, and Vrille were reported to promote the growth and invasiveness of the *Ras^V12^,scrib^-^*tumours (53–57). These data acquired from transcriptomics or RNA-seq have shown the presence of multiple transcription factors in the same genotype suggesting that in these tumour cells several pathways and signals may be simultaneously active that culminate in upregulation of several different transcription factors. These studies also confirm several transcription factors are active based on activation of their target genes. However, if any molecular networks are established remained unclear. Here we present evidences to show that a tumour cell specific network is formed in *Ras^V12^,scrib^-/-^*tumour cells. This network is comprised of Yki, JNK, Wg and Caspases that acts to promote fitness and aggressive growth (Fig. 1).

Of these signals, JNK is very well documented as a modulator of Yki activity in the context of cell competition, compensatory proliferation, regeneration and neoplastic tumours (12,14). JNK is a pivotal stress-responsive kinase that promotes malignant transformation and metastasis of tumours. JNK has also emerged as a key paracrine signal that links apoptosis to carcinogenesis (12). In *scrib^-^* cells, JNK signalling suppresses Yki activity and promotes cell competition (Fig. S1) (12,58). Concomitantly, JNK signalling causes non-cell autonomous propagation of Yki in the cells surrounding *scrib*^-^ clones and promotes compensatory proliferation (12,58). Thus, in *scrib^-^* clones in wild-type background, JNK (Fig. S3B) and Yki (Fig. S3C-D) activities are induced in two distinct cell populations, and wild-type levels of Dronc (Fig. 3E) or Wg (Fig. S3F) are sufficient to support cell competition mediated elimination of *scrib^-^* cells (Fig.1, S1D,G). Work from our lab and others showed that suppression of cell death in *scrib^-^* cells led to increased proliferation, and loss of differentiation but not tumourigenesis (59,60). In contrast, we found that these signalling interactions are significantly altered in *Ras^V12^,scrib^-^* cells as Wg, Dronc, JNK and Yki are all robustly upregulated (Fig. 1, S1) suggesting non-apoptotic tumour growth-promoting role for JNK and Dronc, and growth promoting mitogenic roles for Yki and Wg. Using reporter assays for *dronc* and *wg*, we found that *Ras^V12^,scrib^-^* cells show increased transcription of *dronc* (Fig.1L-L’’) and *wg* (Fig.1M-M’’) which was confirmed by qRT-PCR (Fig.S2). In addition, increased JNK activity using phospho-specific JNK antibody and Yki activity by *diap1-lacZ* reporter were confirmed in the *Ras^V12^,scrib*^-^ Tumour cells (Fig. S1). These findings are consistent with recent reports that JNK signalling activity is converted from anti- to pro-tumour growth through downregulation of Hippo signalling which ultimately leads to Yki activation (14,32). Both Yki/YAP and JNK are linked to Wg signalling, as Wg is induced in a JNK-dependent manner during regenerative growth, tumourigenesis, and compensatory proliferation (41,61,62), and interacts with Yki and the Hippo pathway during organ development and tumourigenesis (63–65). In addition, misregulation of the Hippo, JNK or Wg pathway is also linked to activation of caspases mediated apoptosis, and recently mild caspase induction was shown to promote tumour growth (66,67). Thus, the identification of this molecular network is significant in the context of inter-cellular interactions that promote tumour growth.

An assessment of the roles of JNK, Yki, Dronc and Wg in *Ras^V12^,scrib^-^*tumourigenesis revealed several interesting insights. First, we found that JNK, Yki, Dronc, and Wg were all required for aggressive growth of *Ras^V12^,scrib^-^* induced tumours as downregulation of these signals individually resulted in a significant reduction of tumour growth (Fig.2A). Second, we observed that in the absence of tumour promoting signals *Ras^V12^,scrib^-^* induced tumours show reduced survival and fitness (Fig.2). Third, we identified that the four tumour promoting signals form a tumour cell specific signalling module in which Wg acts upstream of Dronc which, in turn, acts upstream of a JNK-Yki mediated positive feedback signal amplification loop (Fig.3,4). Signal amplification of Yki and JNK activities caused by the JNK-Yki positive feedback loop plays a key role in promoting tumourigenesis in *Ras^V12^,scrib^-^*cells. Fourth, we confirmed that in other instances of oncogenic cooperation (*Ras^V12^//scrib^-^*) this signalling module can be recapitulated (Fig.6). Fifth, upregulation of JNK and Yki in normal epithelial cells with intact polarity is not sufficient to induce this network and tumour growth (Fig.7, S5). Taken together, our studies reveal that increased cellular fitness promoted by a molecular network comprising Wg, Dronc, JNK and Yki may indeed be a mechanism co-opted for aggressive tumour growth. Furthermore, overexpression of pathway components in normal epithelia of wing discs revealed that the levels of overactivation of these signals and the apical-basal polarity context are extremely important as mild hyperplasia can occur by driving the individual transgenes, but robust overgrowth is not seen in cells with intact polarity. Further, for the JNK pathway a threshold is critical as stress-induced cell death occurs both when levels of JNK reach above or below the homeostatic levels.

Another possible mechanism is the differential success of cancer and neighbouring normal cells in competing for survival and other extracellular signals in terms of actively internalizing or limiting the extracellular spread of critical signals. In this context, we observed steep differences in the non-cell autonomous spread of Yki activity between *scrib^-^* (Fig.5), *Ras^V12^ scrib^-^* (Fig.S1C,C’), and *en>Yki; scrib^-^* (Fig.5C) cells. Non-cell autonomous Yki activity spreads to 3-5 cells in *scrib^-^* cells, several cells (8–10) in *en>Yki; scrib^-^* and only a few cells (∼2) around the *Ras^V12^,scrib^-^* clones. Thus, limiting the extracellular spread of key signalling components like Yki or Wg may impact the growth potential of cancer cells. The mechanisms by which cancer cells limit the spread of these signals should be elucidated in future studies.

The biological significance of our findings is validated by other studies that show formation of context-dependent Yki/YAP-mediated signalling loops as a *bona fide* mechanism for tumour growth (68,69). YAP is not only regulated by signalling pathways like Wnt, TGFβ and Notch, but YAP can also collaborate with key cancer pathways to form transcriptional complexes that alter transcriptional programs specifically in cancer cells (70,71). At least three different mechanisms have been reported. First, YAP and its cognate transcription factors like TEAD control transcription by binding to promoters of target genes. Second, YAP collaborates with other transcription regulators to alter gene expression, for example, in colon cancer cell lines YAP1, β-catenin and TBX5 transcriptional complex regulate (TCF- or TEAD-independent) target genes (72). Third, YAP/TEAD bind distal regulatory elements to regulate transcription of target genes (73–75). ChIP-seq and deep-sequencing studies in cancer and normal cells showed that several YAP-binding regions also show a consensus motif for AP-1 transcriptional factors suggesting that YAP/TEAD and AP-1 cooperatively regulate target genes representing a cross-talk between YAP and JNK signalling (74,75).

These findings are especially interesting in light of our identification of the tumour-cell-specific Wg-Dronc-JNK-Yki molecular network and the JNK-Yki positive feed-back signal amplification loop in the *Ras^V12^ scrib^-^* cells. Like YAP, *Drosophila* Yki is known to regulate transcription by binding to promoter regions of genes (76). Similarly, Yki can form transcriptional complexes that cooperatively alter gene expression in cancer cells, for example, Yki and the Ecdysone receptor coactivator Taiman were shown to alter transcriptional output of Yki inducible Taiman dependent genes in cells with hyperactive Yki, and neoplastic tumour growth (77). Based on our data, it is possible that under conditions of oncogenic cooperation, Yki and the *Drosophila* AP-1 transcription factors may (cooperatively) bind other regulatory regions to drive altered transcriptional output in cancer cells. This conclusion is further strengthened by the finding that dFos, a component of the *Drosophila* AP-1 (*Drosophila* Jun and Fos heterodimer) transcription factor, is required for the JNK-mediated invasiveness of *Ras^V12^, scrib^-^* tumours (35,55). Our data is significant in light of emerging data from cancer genome studies that revealed that somatic mutations accrued by precancerous cells essentially activate hallmark cancer pathways that converge on a small number of protein complexes and signalling cascades (6). Our genetic analyses indicate the hierarchy of one such signalling interaction with Wg acting upstream of Dronc, JNK and Yki, and in the future, it will be interesting to identify the molecular mechanisms underlying the establishment and regulation of this module. Our approach of modelling altered signalling interactions in a genetically tractable model is a powerful way to reconstruct key biologically meaningful changes in signalling pathways in cancer cells and provides a functional framework to study the changes in signalling pathway interactions that will generate new insights on mechanisms that promote tumour growth and progression.

## Author Contributions

Conceptualization, M.K-S and A.S.; methodology, M.K.S. and I.W.; formal analysis, I.W., K.G., A.R., M.K-S; investigation, I.W., M.K.S. and A.S.; resources, M.K-S and A.S.; writing, original draft preparation, I.W., M.K-S.; writing, review and editing, I.W., A.R., M.K-S., A.S.; supervision, M.K-S.; project administration, M.K-S; funding acquisition, M.K-S., A.S.

## Institutional Review Board Statement

Not applicable.

## Informed Consent Statement

Not applicable.

## Acknowledgments

We would like to thank Profs. A. Bergmann, G. Halder, B. Hay, K. D. Irvine, H.D. Ryoo and Tin Tin Su, and the Bloomington *Drosophila* Stock Center, the *Drosophila* Genetics Resource Center (Japan), and the Developmental

Studies Hybridoma Bank for flies and antibodies; and the members of the Kango-Singh and Singh Labs for comments and suggestions on the manuscript. IW, KG and AR acknowledge the Graduate School at University of Dayton for the Graduate Student Summer Fellowships.

## Supplementary Figures

**Fig. S1 (related to** Fig. 1**): Growth of scrib mutant clones and the role of Yki and JNK signalling in promoting *Ras^V12^ scrib^-^* clones**. (A-B’) Panels show activation of Casp 3* (A,B red, A’B’ grey) in *scrib^-^* (A GFP) and *scrib^-^,P35* (B GFP) MARCM clones. (C-F) Show expressing of *diap1-lacZ* (using anti-β-gal antibodies) in wild-type (C), flp-out clones of Yki (D) and in MARCM clones of *Ras^V12^ scrib^-^* (E red, E’ grey). The dot plot in F depicts the pixel intensity from indicated genotypes for expression of diap1-lacZ. (G-I) MARCM clones (GFP, green) of wild-type (G) and *Ras^V12^ scrib^-^*(H) genotypes stained for pJNK (G, H red, G’, H’ grey). The dot plot in I show changes in pJNK levels for the tested genotypes. For statistical analysis in F and I, a student’s t-test was used to determine significant difference (n=5, p≤ 0.05). The dashed yellow line in panels E, E’, H, H’ marks the clone boundary. The magnification and orientation of discs is identical in all panels.

**Fig. S2 (related to** Fig. 1**): Upregulation of *wg* and *dronc* expression in *Ras^V12^ scrib^-^* clones.** (A, B) graphs show fold-change in expression of wg (A) and dronc (B) in the indicated genotypes assessed by qRT-PCR.

**Fig. S3 (related to** Fig. 2**): Assessment of cell survival in *Ras^V12^ scrib^-^* clones.** Panels show a comparison of expression of DIAP1 (red, grey) in wild-type, *scrib^-^* and *Ras^V12^ scrib^-^* MARCM clones (green). Yellow arrowheads indicate downregulation of DIAP1 levels in *scrib^-^* clones (middle row). The magnification and orientation of discs is identical in all panels.

**Fig. S4 (related to** Fig. 2**): Effects of Downregulation of network components in wild-type cells.** Panels show MARCM clones of wild-type cells in which we depleted (A-B’”) Wg (*UASgg^S9a^;<+/<+*), (C-D’”) Dronc (*UASDronc^RNAi^; <+/<+*), (E-F’”) JNK (*UASBsk^DN^; <+/<+*) or Yki (*UASYki^N+CRNAi^; <+/<+*) and tested effects on expression of DIAP1 (A, C, E, G red, A”, C”, E”, G” grey), DCP1 (A, C, E, G blue, A”’, C”’, E”’, G”’ grey), Wg (B, D, F, H red, B”, D”, F”, H” grey), and pJNK (B, D, F, H blue, B”’, D”’, F”’, H”’ grey). The magnification and orientation of discs is identical in all panels.

**Fig. S5 (related to** Fig. 5**): Effects of loss of *scrib* on the molecular network.** Panels show *scrib^-^* MARCM clones (green) in eye imaginal discs stained to assess expression of pJNK (A, B red, A’ B’ grey), *diap1-lacZ* using anti-β-gal antibodies (C red, C’ grey), Yki (D red, D’ grey), Dronc (E red, E’ grey) and Wg (F red, F’ grey). The magnification and orientation of discs is identical in all panels.

**Fig. S6 (related to** Fig. 7**): Activation of individual network components in polarity intact cells cause mild effects.** Panels show wing imaginal discs from (A-B”) *MS1096>UAS Yki*, (C-D”) *MS1096>UASproDronc*, (E-F”) *MS1096>UASjun^aspv^,* and (G-H”) *MS1096>UASArm^S10^* stained for DIAP1 (A, C, E, G red, A’, C’, E’, G’ grey), DCP1 (A, C, E, G blue, A”, C”, E”, G” grey), Wg (B, D, F, H red, B’, D’, F’, H’ grey) and pJNK (B, D, F, H blue, B”, D”, F”, H” grey). The magnification and orientation of discs is identical in all panels.

