## Supplementary figures and images for "A Tumour-Specific Molecular Network Promotes Tumour Growth in *Drosophila* by Enforcing a JNK-YKI Feedforward Loop"

### Supplementary Figure 1

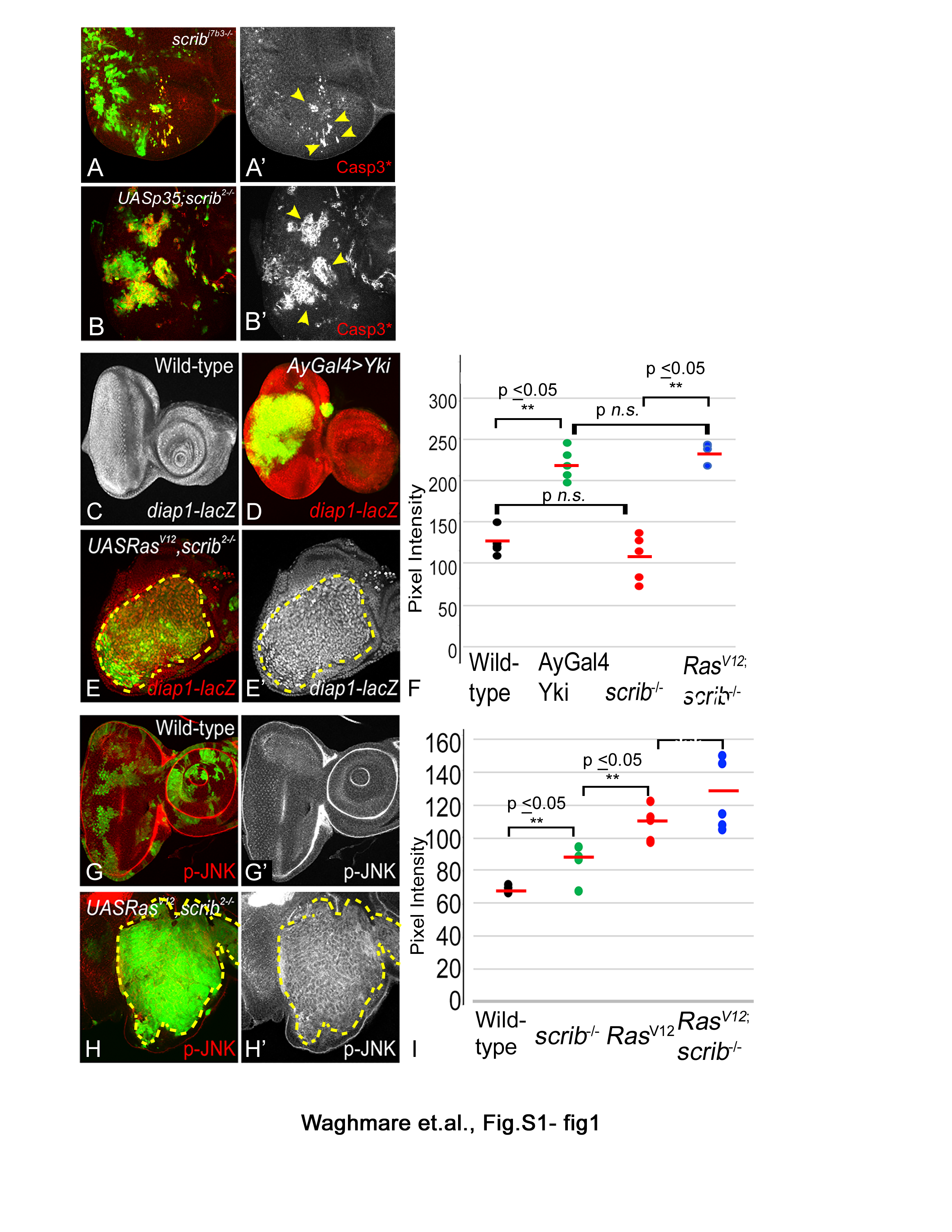

### Supplementary Figure 2

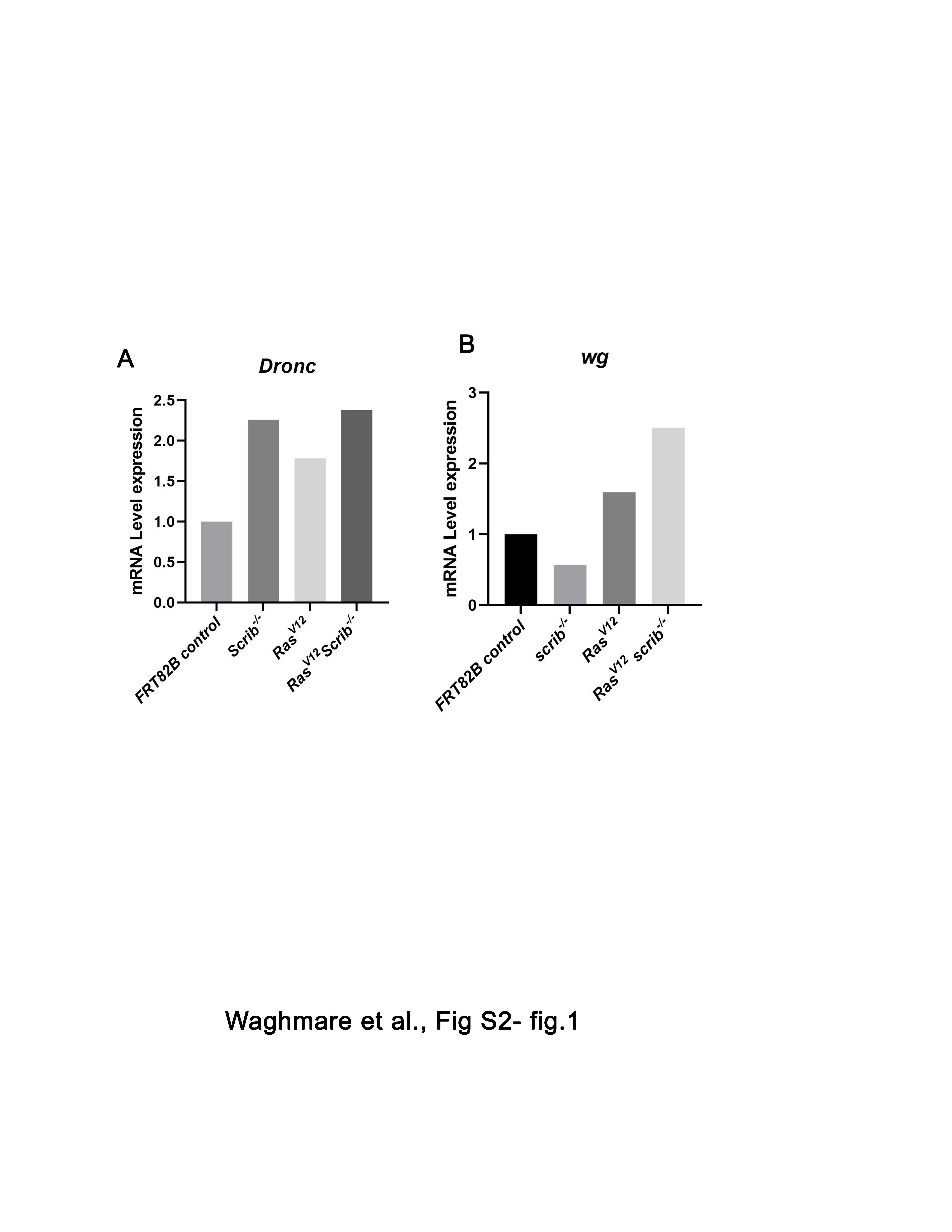

### Supplementary Figure 3

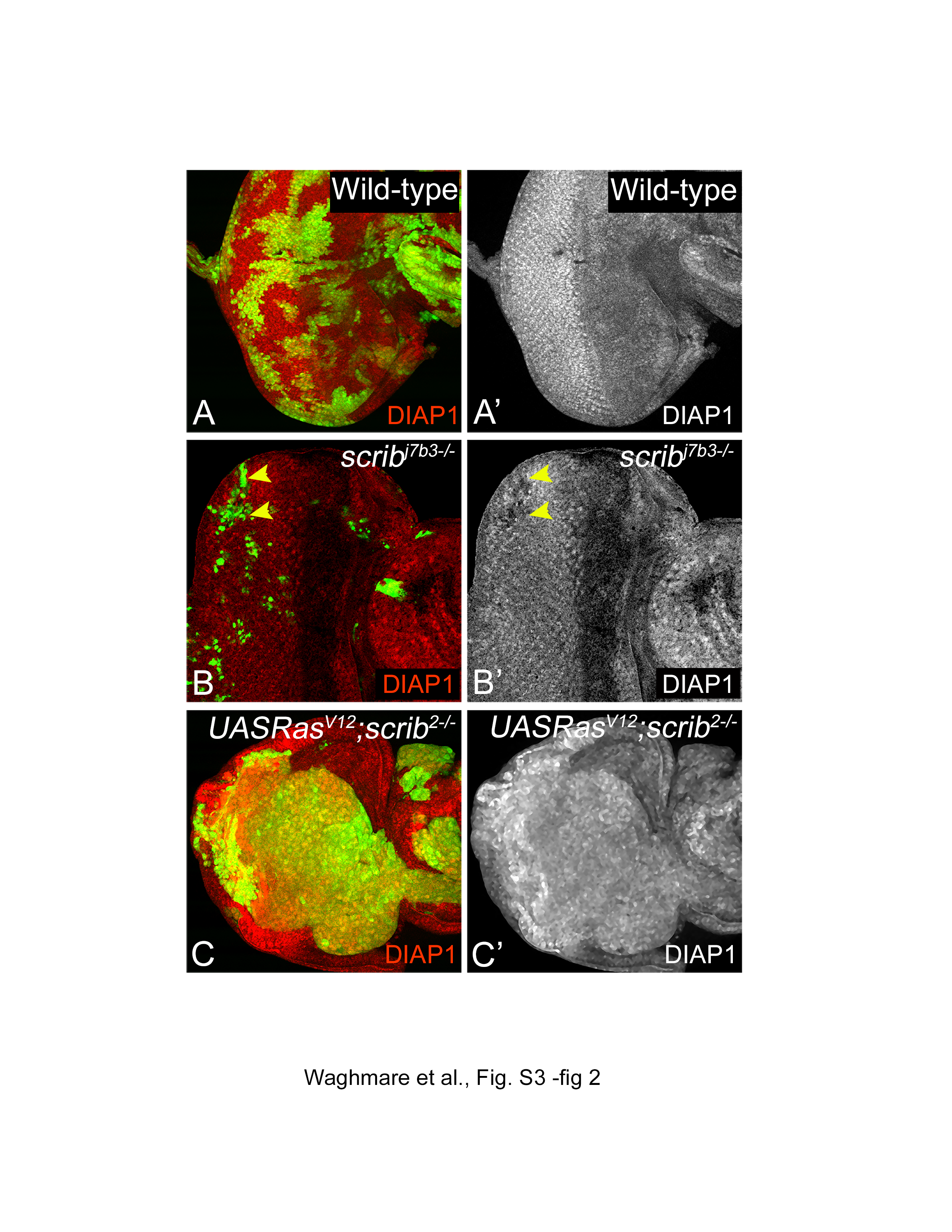

### Supplementary FIgure 4

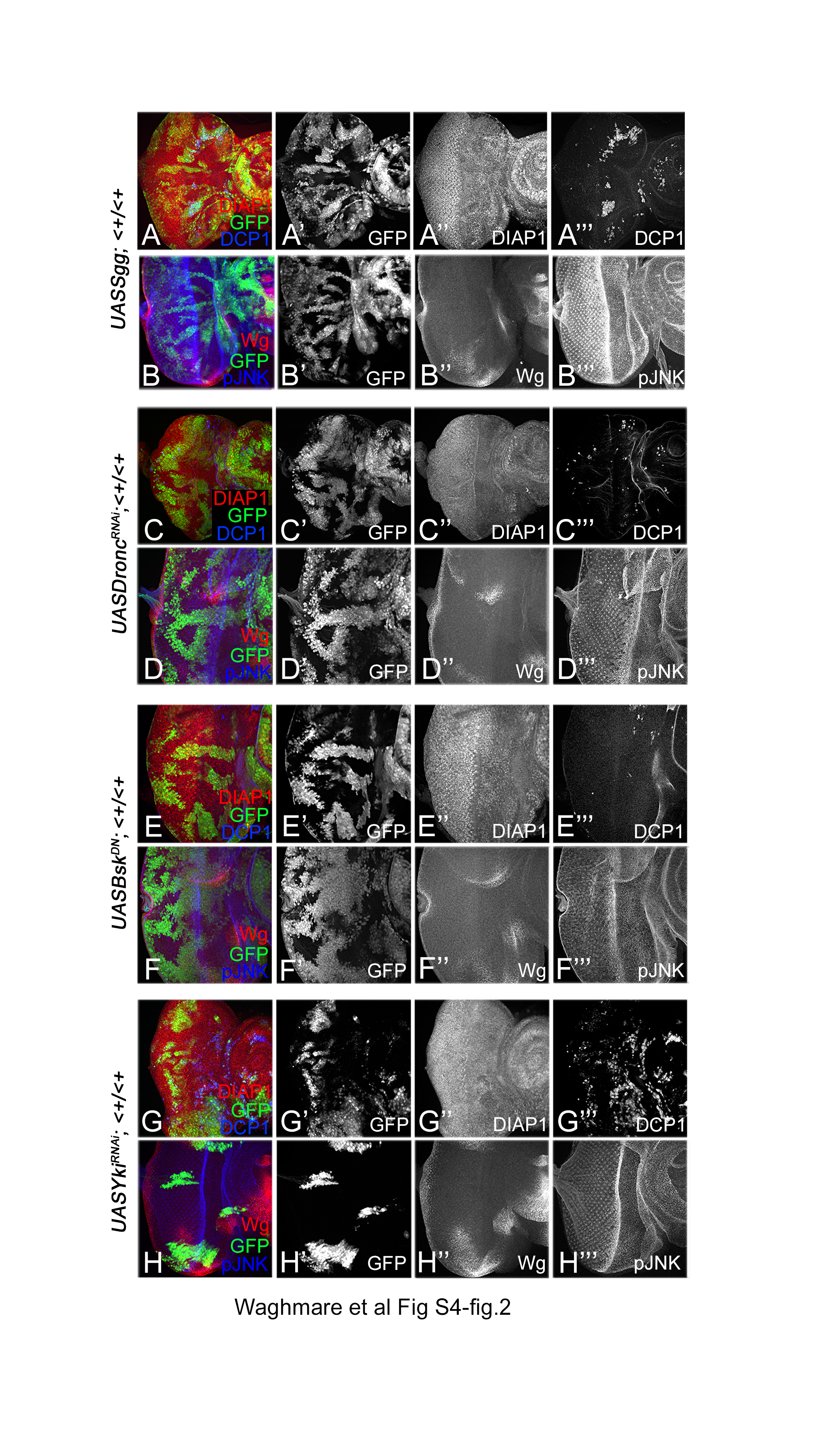

### Supplementary Figure 5

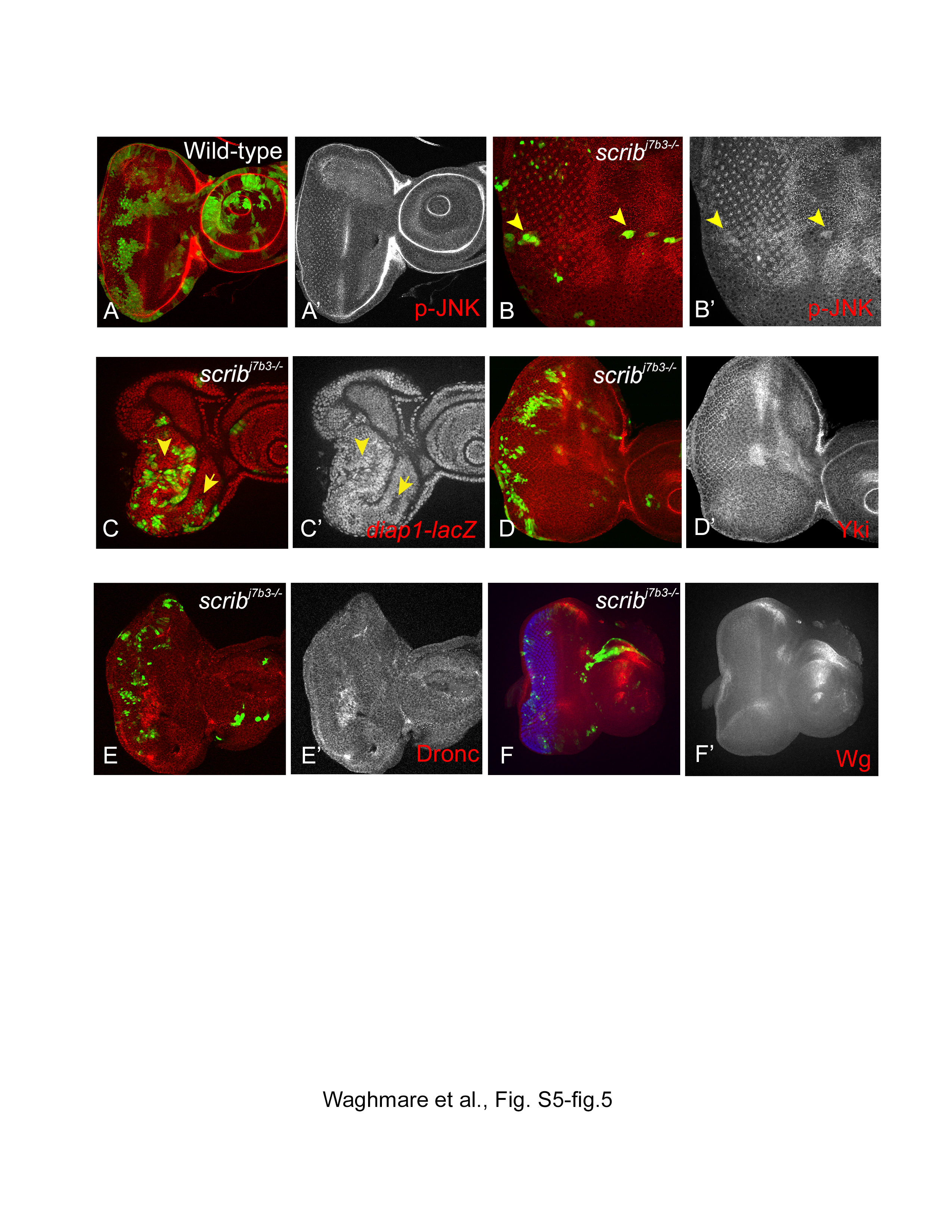

### Supplementary Figure 6

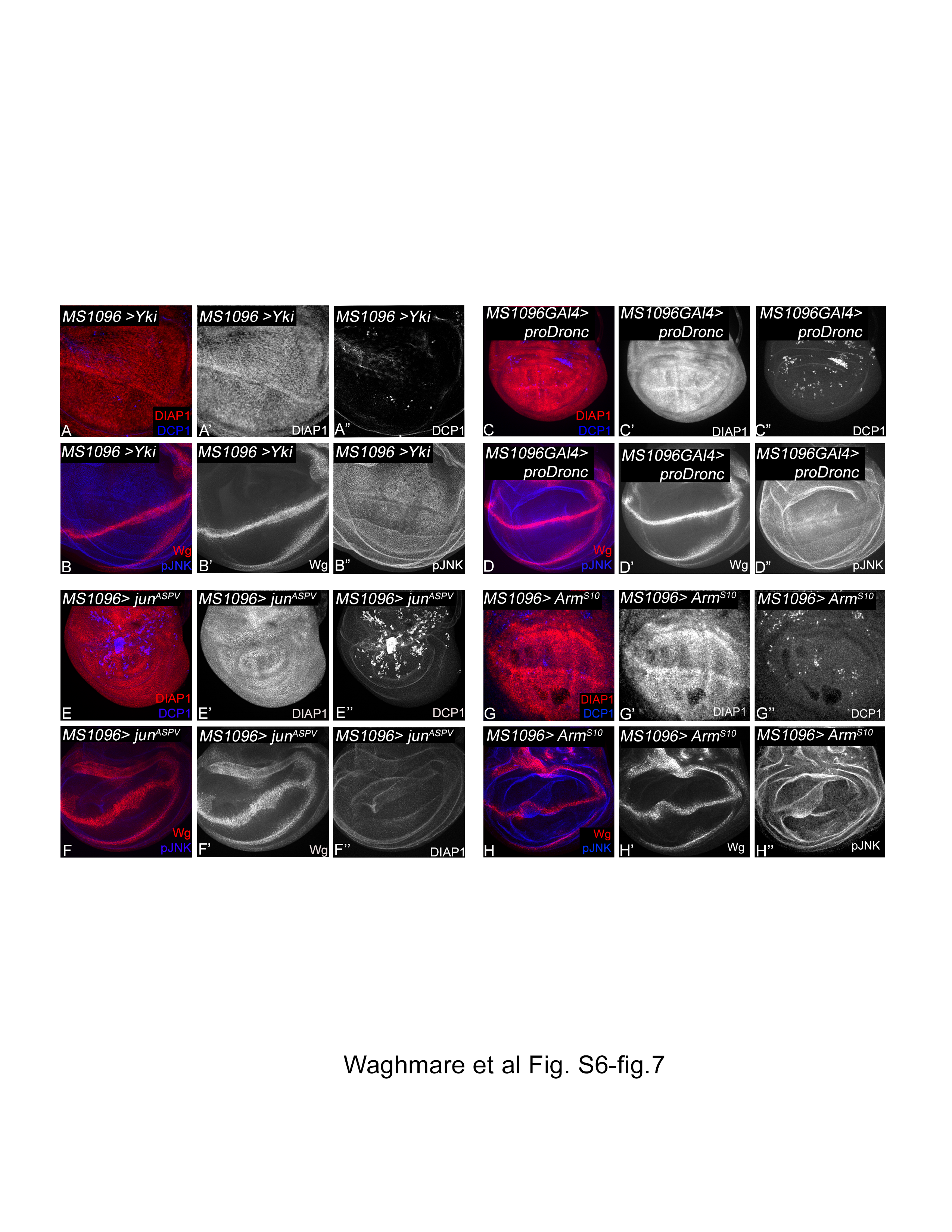
